# PathoFact 2.0: An Integrative Pipeline for Predicting Antimicrobial Resistance Genes, Virulence Factors, Toxins and Biosynthetic Gene Clusters in Metagenomes

**DOI:** 10.1101/2024.12.09.627531

**Authors:** Júlia Ortís Sunyer, Luis F. Delgado, Oskar Hickl, Cedric C. Laczny, Patrick May, Paul Wilmes

## Abstract

**Summary:** Antimicrobial resistance genes (ARGs) and virulence factors (VFs) are central contributors to the global health crisis surrounding drug-resistant infections. PathoFact, a bioinformatics pipeline introduced in 2021, provides insights into ARGs, VFs, and bacterial toxins from metagenomic data. However, recent advancements in bioinformatics highlight the need for an updated version of PathoFact. We introduce PathoFact 2.0, an enhanced pipeline for improved ARG, VF, and toxin prediction. Key updates include an updated machine learning (ML) model for VF identification, a new ML model for toxin identification, expanded hidden Markov model profiles, and the antiSMASH 7.0 integration for predicting biosynthetic gene clusters. These upgrades make PathoFact 2.0 a more powerful, user-friendly platform for predicting microbiome-based pathogenicity and resistance, offering a crucial tool for better understanding and addressing the challenges posed by antimicrobial resistance and infectious diseases.

**Availability and Implementation:** PathoFact 2.0 is available for download at https://gitlab.lcsb.uni.lu/ESB/PathoFact2/.

## 1. Introduction

Microbiomes are complex communities of bacteria, archaea, microeukaryotes, and viruses within human, animal, and environmental habitats. They harbour commensal and pathogenic microorganisms. These communities contribute to infectious disease emergence and antibiotic resistance by serving as reservoirs for antimicrobial resistance genes (ARGs), virulence factors (VFs), and toxins (Inda-Díaz et al., 2023).

ARGs provide resistance to antimicrobials through gene mutations or horizontal gene transfer via mobile genetic elements (MGEs) like plasmids and transposons (Michaelis and Grohmann, 2023). VFs enable pathogens to colonize and survive in hosts, acting as cytosolic, membrane-associated, or secretory components, which support metabolism, immune evasion, and host defence interference. (Sharma et al., 2017). VFs are also transferable across microbial populations via MGEs, spreading virulence (Blair et al., 2015; Rodríguez-Beltrán et al., 2021).

Bacterial toxins, including exotoxins and endotoxins, disrupt host processes and manipulate immune responses. Some toxins impair protein synthesis, destroy blood cells, or affect the nervous system. They are key VFs that promote bacterial colonization and infection, as seen in cholera caused by *Vibrio cholerae* (Conner et al., 2016).

Biosynthetic gene clusters (BGCs) drive the synthesis of metabolites that may influence pathogenicity; for example, *P. aeruginosa* and *Burkholderia* species produce siderophores, rhamnolipids, toxoflavin and phenazines, enhancing virulence (Elshafie and Camele, 2021; Lau et al., 2004; Lybbert et al., 2020).

ARGs, VFs, and toxins have a profound impact on human health. The United Nations has declared antimicrobial resistance a global threat, with drug-resistant infections currently causing an estimated 1.27 million deaths annually—a figure projected to reach 10 million by 2050 if unaddressed (UN Environment Programme, 2024a, 2024b). Accurate prediction of ARG and VF profiles is therefore critical for anticipating infection severity, optimizing treatment strategies, and reducing mortality from pathogenic infections.

Predicting and annotating VFs, toxins, and ARGs is challenging due to limited well-annotated data (Bansal et al., 2022), complex mechanisms involving gene transfer, mutations, and multifactorial interactions. Traditional annotation methods relying on sequence similarity may overlook novel VFs and ARGs, while machine learning offers robust solutions through pattern recognition, allowing for accurate predictions despite limited training data. An integrated bioinformatics pipeline further enhances analysis by examining VFs, ARGs, toxins, signal peptides, and BGCs, improving insights into pathogenicity and resistance, streamlining workflows, and simplifying data interpretation.

PathoFact, introduced in 2021, provides a comprehensive pipeline for predicting ARGs, VFs, and bacterial toxins from metagenomic data (de Nies et al., 2021). Despite the development of several other tools for predicting these features individually (Dong et al., 2024; Rathore et al., 2024) only HyperVR (Ji et al., 2023), has attempted simultaneous prediction. However, HyperVR’s repository is no longer available online, and the Zenodo archive from its original submission lacks the necessary databases.

Here, we present PathoFact 2.0, which improves existing functionality and extends PathoFact’s scope. PathoFact 2.0 supports protein sequences and contigs as input. It features updates to ARG, VF, and toxin prediction, improved ML models for VFs, new ML models for toxins and an updated hidden Markov model (HMM) database. Additionally, PathoFact 2.0 introduces the BGCs prediction. This new feature complements ARG, VF, and toxin prediction capabilities.

## 2. Methods

### Pipeline Installation and Implementation

PathoFact 2.0 introduces updates to the ML algorithms and integrated tools, and it simplifies the installation process. It has been developed using Python (version 3.12) (RRID:SCR_024202) and Snakemake (version 7.25.0) (Köster and Rahmann, 2012). PathoFact 2.0 is open-source (with license GNU General License v3.0 or later) and accessible at https://gitlab.lcsb.uni.lu/ESB/PathoFact2.

### Additional Functionalities

PathoFact 2.0 integrates SignalP (version 6) (Teufel et al., 2022) and antiSMASH (version 7.0) (Blin et al., 2023), both optional features to accommodate diverse research needs. SignalP is designed to predict the presence and location of signal peptides in protein sequences. It requires a separate license and must be requested. AntiSMASH is designed for the identification and annotation of BGCs in bacterial and fungal genomes.

### Pipeline Structure

Unlike version 1.0, which only supports contigs, PathoFact 2.0 handles both nucleotide sequence of contigs and protein sequence FASTA files as input, with proteins dereplicated to retain non-redundant sequences (100% identity and coverage). For contig-based inputs, open reading frames are predicted using Pyrodigal-gv (version 0.3.2) (Larralde, 2024), followed by the detection of MGEs and phages using geNomad (version 1.8.0) (Camargo et al., 2024). GeNomad only processes nucleotide sequences; hence, MGEs and phages are not detected in protein sequence inputs. Based on user configuration, the pipeline then analyzes the processed sequences using BGC, ARG, VF, and toxin prediction modules. The information is compiled into individual module reports and a comprehensive final report, incorporating details from SignalP and geNomad. Additionally, PathoFact 2.0 generates a FASTA file of proteins identified as VF, ARG, or toxin related.

### ARG Prediction Updates

ARG prediction in PathoFact 2.0 integrates DeepARG (version 1.0.2) (Arango-Argoty et al., 2018), RGI (version 6) (Alcock et al., 2023), and AMRFinderPlus (version 3.12.8) (Feldgarden et al., 2021). While DeepARG and RGI have been updated from PathoFact 1.0, AMRFinderPlus has been incorporated. Each tool has distinct strengths: DeepARG offers high precision and recall, RGI provides robust predictions with extensive database support, and AMRFinderPlus efficiently identifies resistance genes and mutations using NCBI resources (RRID:SCR_006472). Results include protein IDs, ARG classes, prediction probabilities, database accession numbers, and optional data on signal peptides, plasmids, and virus markers.

### Toxin Prediction Updates

The toxin prediction module now employs a ML model instead of a bit score threshold, enhancing detection accuracy (Table 1). Curated training data was obtained from SwissProt (The UniProt Consortium, 2023), filtered for bacterial, archaeal, fungal and viral toxin sequences, supplemented with entries from toxin-specific databases like T3DB (Wishart et al., 2015), DBETH (Chakraborty et al., 2012), TADB (Shao et al., 2011), SecReT6 (Zhang et al., 2023), and PAT (Liu et al., 2023). MMseqs2 (version 15.6f452) (Steinegger and Söding, 2017) was used to dereplicate the dataset of 1,112,357 protein sequences, yielding 213,363 unique protein sequences in the positive dataset. HMM profiles were built using the Conserved Domain Database (Wang et al., 2023) and selected for annotation if they had a bit score >25.

**Table 1.** Detailed results on the validation of ARG, VF and toxin prediction.

| Dataset |  | Accuracy | Precision | Recall | F1 score |
| --- | --- | --- | --- | --- | --- |
| ARG | HMD-ARG + control VF Test dataset | 83.64 | 99.98 | 75.73 | 86.18 |
| VF | VF + control VF Test dataset | 99.01 | 99.72 | 97.93 | 98.82 |
| Toxin | Toxin + control Toxin Test Dataset | 96.73 | 99.78 | 93.65 | 96.62 |

The negative protein dataset (i.e., proteins that are not ARGs, VF or toxins) for the ML models was constructed by selecting SwissProt sequences lacking ARG, VF, and toxin keywords (KW-0568, KW-0843, KW-0800, KW-0046, KW-9995) and restricted to bacteria, archaea, fungi, and viruses. Additionally, proteins from non-pathogenic organisms to humans (Supplementary Table 1) were included from NCBI. Potential ARGs, VFs (with high probability), and toxins were filtered out based on PathoFact 1.0 predictions, resulting in 213,130 non-redundant (100% identity and coverage), protein sequences. The ML model training set comprised 80% of the positive and negative datasets, with 20% reserved for testing.

Synthetic Minority Oversampling Technique (SMOTE) was used to tackle the datasets imbalance (Chawla et al., 2002). Several ML models were build using Sckit-learn (version 1.5.2) (Pedregosa et al., 2011) and tested [Random Forest (RF) and XGBoost, with k-mers 3 to 8 or protein sequence composition features (amino acid composition, dipeptide composition, composition, transition and distribution) as features]. The best-performing model used a RF with 5-mer features (default hyperparameter setting) with a probability threshold of 0.8 for classification as a toxin. The toxin prediction module generates a report containing the proteinID, protein domains, bitscore, toxin ML probability, other identical proteins found in the sample, and optionally SignalP, plasmid marker, and virus marker information.

### VF Prediction Updates

The VF prediction model was refined and updated with new HMM profiles. Training data was derived from SwissProt (The UniProt Consortium, 2023), selecting sequences annotated with virulence keyword [KW-0843] and expanded using the Virulence Factor Database (VFDB) (Liu et al., 2022). After dereplication (100% identity and coverage), the original set of 32,511 sequences using MMseqs2, the dataset comprised 30,695 non-redundant sequences (i.e., positive dataset). We performed a search of the created VF dataset against the CDD HMM profiles, and those with a bit score of 25 and higher were selected as VF HMM profiles for PathoFact 2.0. The HMM profile dataset annotates the predicted VF domains instead of using them as input for the classification as the previous version did.

To create the negative dataset for the ML VF model, we filtered out any possible VFs (with high and low probabilities), ARGs, and toxins based on PathoFact 1.0 predictions from the previously described negative dataset. This resulted in a dataset consisting of 41,774 protein sequences. 80% of the sequences in the positive and negative datasets were used as the training set, while 20% were part of the test set for the ML models. SMOTE was applied to address the imbalance in the datasets. Several ML models [RF and XGBoost with k-mers 3 to 8 or protein sequence composition features (amino acid composition, dipeptide composition, composition, transition and distribution) as features] were tested. The best-performing model used XGBoost with protein sequence composition features (learning_rate’: 0.1, ‘max_depth’: None, ‘n_estimators’: 2000) with a probability threshold of 0.9 for classification as a VF. This module generates a report containing the proteinID, protein domains, bitscore, Virulence factor ML probability, other identical proteins found in the sample, and optionally SignalP, plasmid marker, and virus marker information.

## 3. Results

### PathoFact 2.0 pipeline performance evaluation

To assess the performance of PathoFact 2.0, we conducted validations for predicting VFs, ARGs, and toxins. For ARG prediction assessment, we used the HMD-ARG database (Li et al., 2021) and applied MMseqs2 at 100% identity and coverage to remove redundant sequences, resulting in a final set of 17,282 unique protein sequences. Test datasets previously described here were used to evaluate the model performance on VF and toxin predictions. The quality of these predictions was evaluated using the precision, recall, F1 score, and accuracy metrics (Table 1).

To evaluate PathoFact 2.0 at the contig level, we analyzed publicly available complete genomes from pathogenic and non-pathogenic bacteria, including various *Escherichia coli* strains (Figure 1, Figure S1 and Table S2). Comparative analysis was conducted between PathoFact 2.0, PathoFact 1.0, and metaVF, a toolkit identifying species-level VFs associated with pathobionts (Dong et al., 2024).

**Figure 1.**
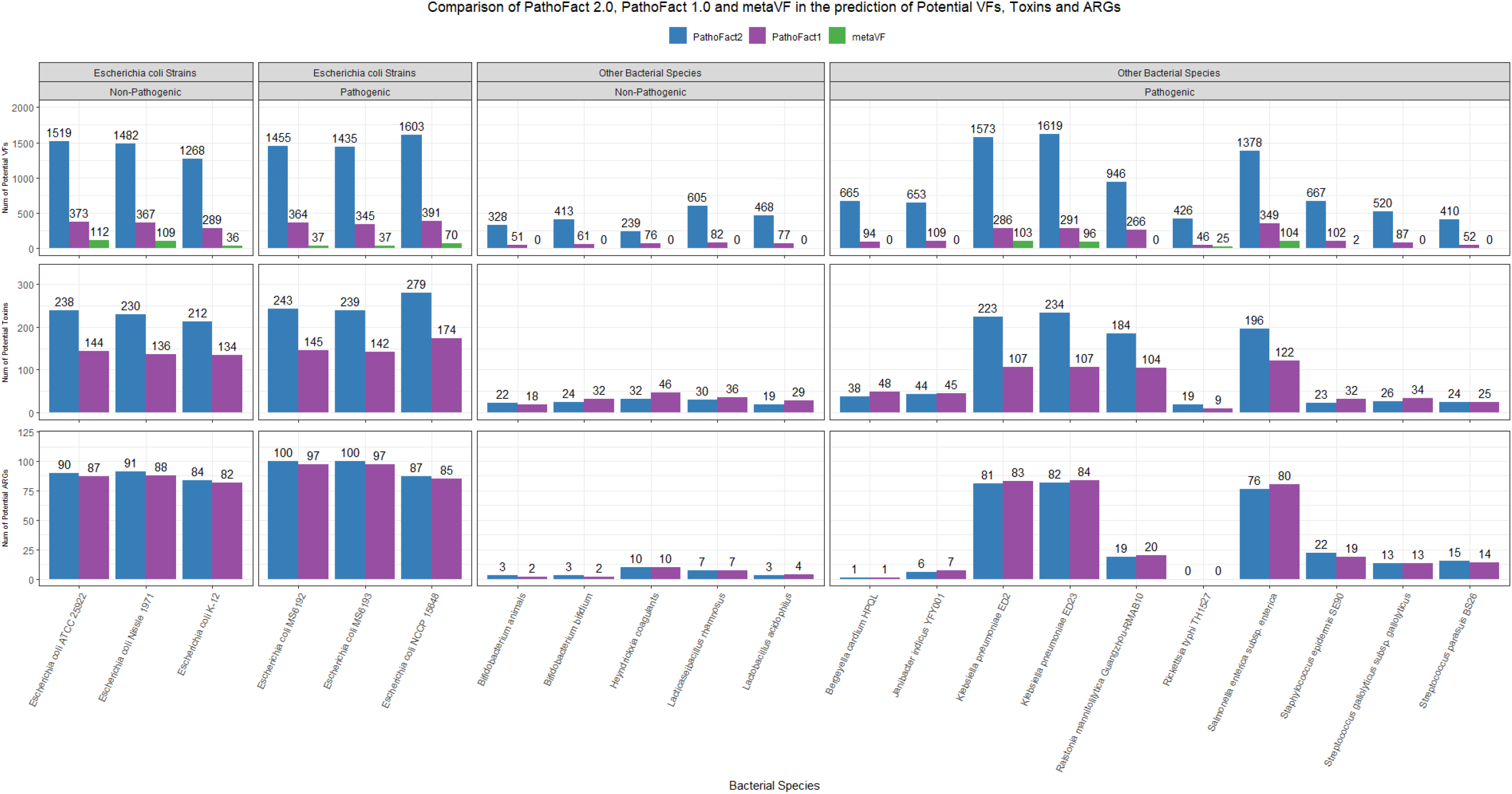
Comparison of PathoFact v2.0 (in blue), metaVF (in green) and PathoFact v1.0 (in purple) in the prediction of potential virulence factors (VFs), toxins and antimicrobial resistance genes (ARGs).

PathoFact 2.0 predicted more VFs and toxins than PathoFact 1.0 and metaVF (Figure 1), and more VF, ARGs and toxins encoded in MGEs (Figure S1). MetaVF failed to identify any virulence factors in pathogenic strains (Figure 1) and struggled to predict VF found in plasmids where they have been reported (Figure S1). Furthermore, PathoFact 2.0 can identify BGCs, some of them are also classified as VF or toxins (Table S2).

The number of VF predictions identified by PathoFact 2.0 was compared to VFs documented in established databases, focusing on three well-characterized pathogenic species. We used MMseqs2 (100% identity and coverage) to dereplicate VFs from three datasets: VFDB setB (Liu et al., 2022) and eVFGC and pVFGC from metaVF (Dong et al., 2024). The eVFGC set includes 62,332 VF gene orthologues and alleles across 135 bacterial species, representing 18,521 genomes and 3,559 species, while pVFGC, a subset of eVFGC, focuses on pathogenic VF gene alleles only. This resulted in the following non-redundant VF counts: for *Salmonella enterica*, 3,190 (eVFGC), 829 (pVFGC), and 1,442 (VFDB setB); for *Escherichia coli*, 4,536 (eVFGC), 1,578 (pVFGC), and 2,028 (VFDB setB); for *Klebsiella pneumoniae*, 2,548 (eVFGC), 740 (pVFGC), and 595 (VFDB setB). These counts align closely with those predicted by PathoFact 2.0 (Figure 1), supporting the validity of PathoFact 2.0 predictions.

The dual roles of certain proteins, such as multidrug efflux pumps and β-lactamases, as ARG and VF elements are well-documented (Beceiro et al., 2013) and pose challenges for precise functional classification. Additionally, some non-pathogenic strains contain VF and toxin genes (Niu et al., 2013), indicating that PathoFact should primarily be used as a preliminary screening tool. Genes identified as potential candidates by PathoFact require further comparative analyses and experimental validation to confirm their pathogenic potential.

PathoFact 2.0 significantly extends the first version capabilities, providing a robust and efficient platform for the comprehensive identification of VFs, ARGs, toxins, and BGCs while incorporating MGE detection and enhancing overall prediction quality.

## Acknowledgements

The experiments presented in this paper were carried out using the HPC facilities of the University of Luxembourg (Varrette et al., 2022). The manuscript also passed the Luxembourg Centre for Systems Biomedicine internal pre-publication check designed to ensure FAIRness and reproducibility.

## Supplementary Information

**Table S1.**
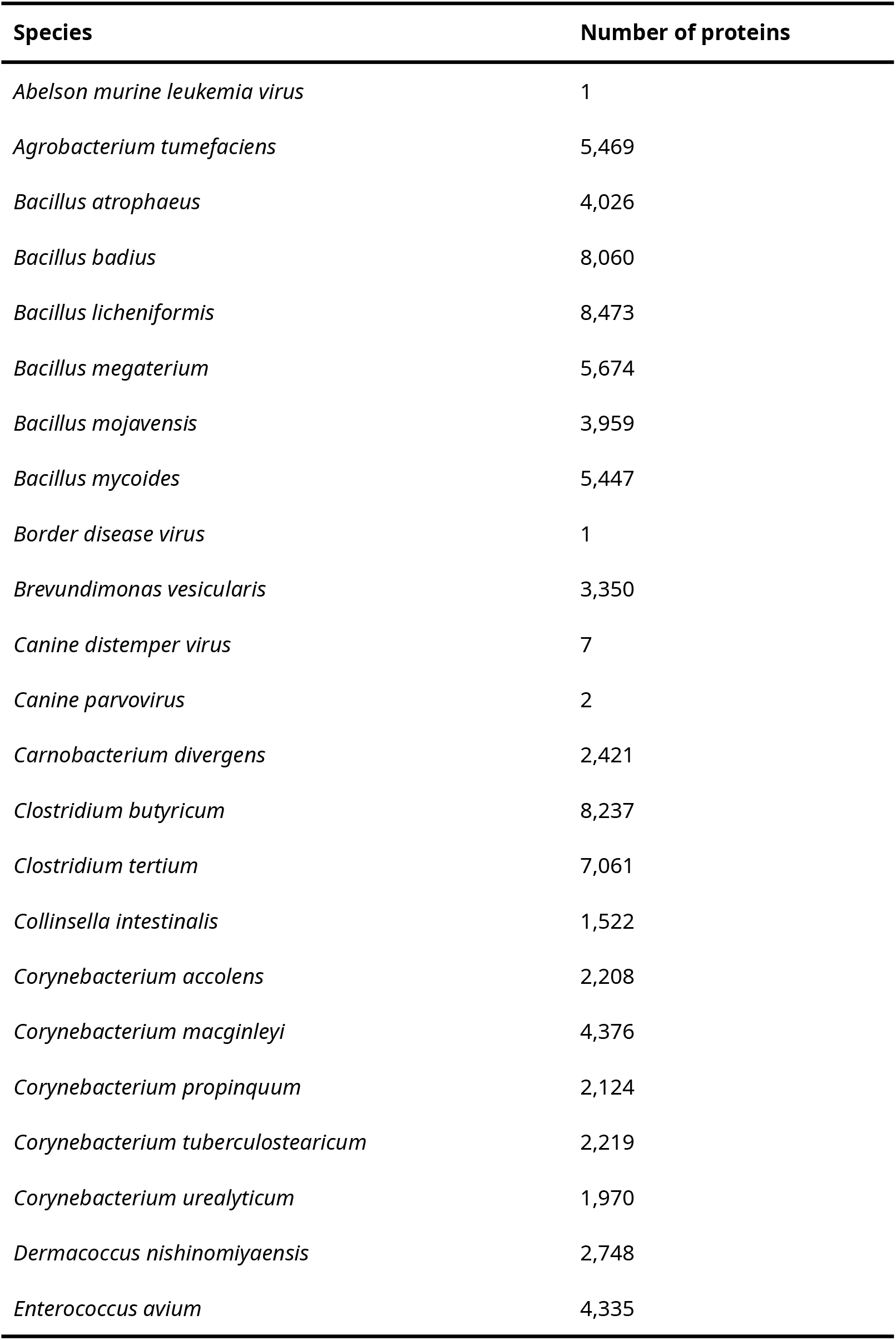
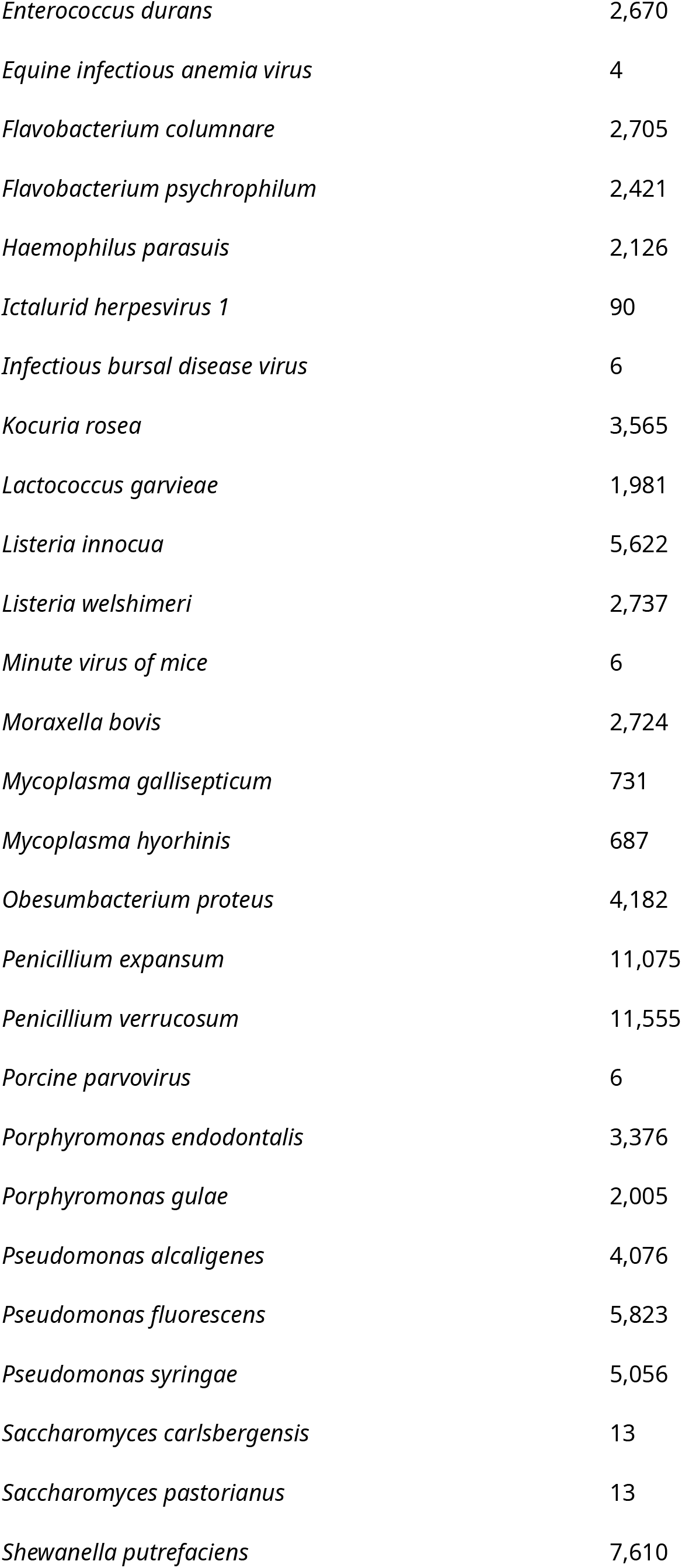
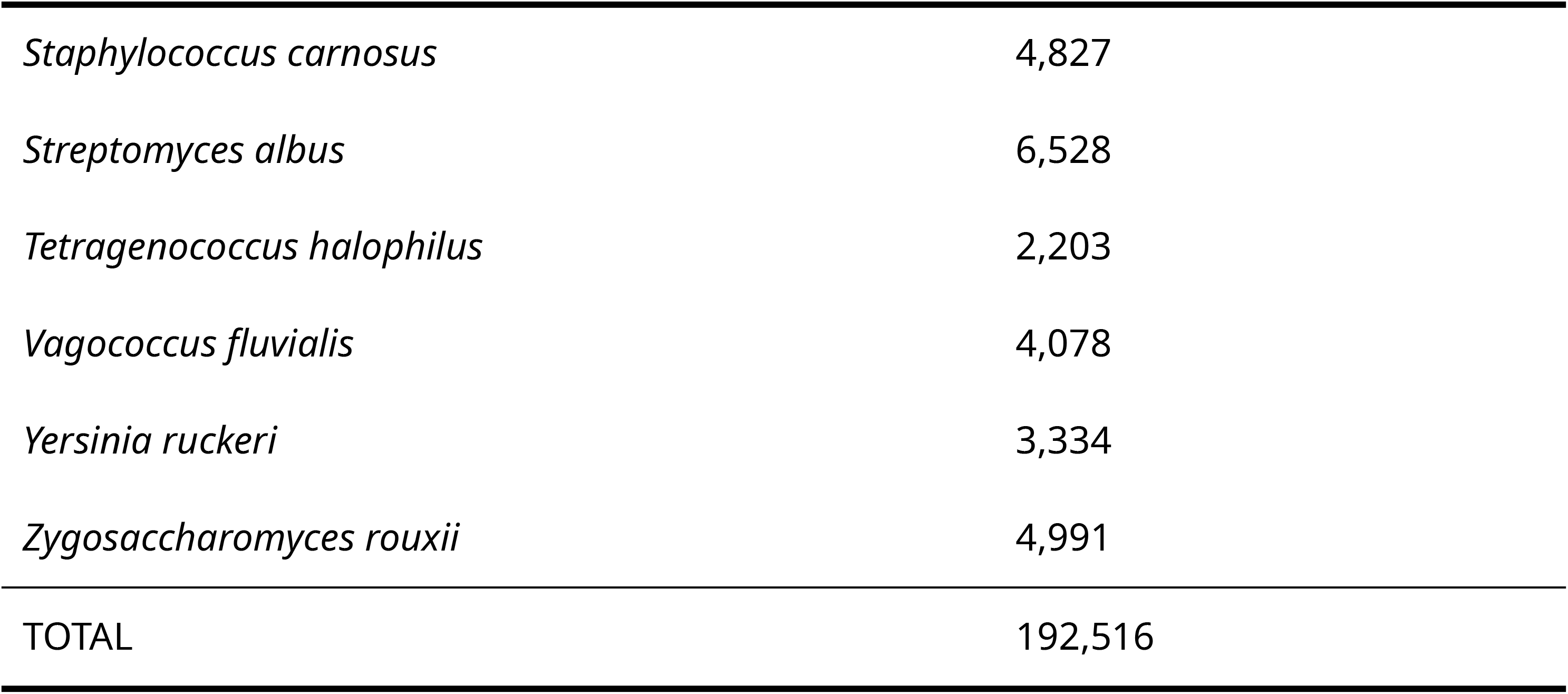
List of microorganisms non-pathogenic to humans and their total protein count obtained from the NCBI Database.

**Table S2.**
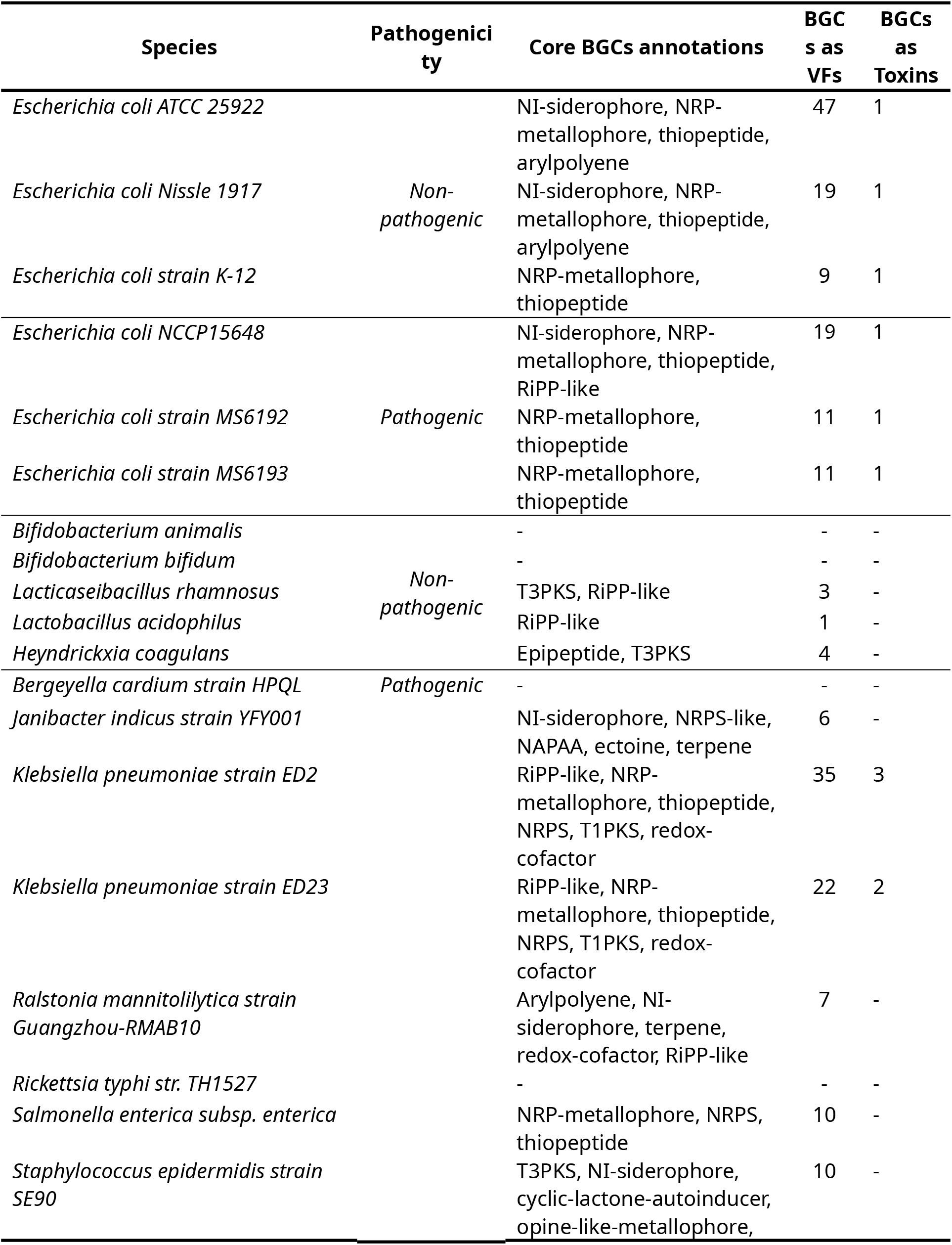
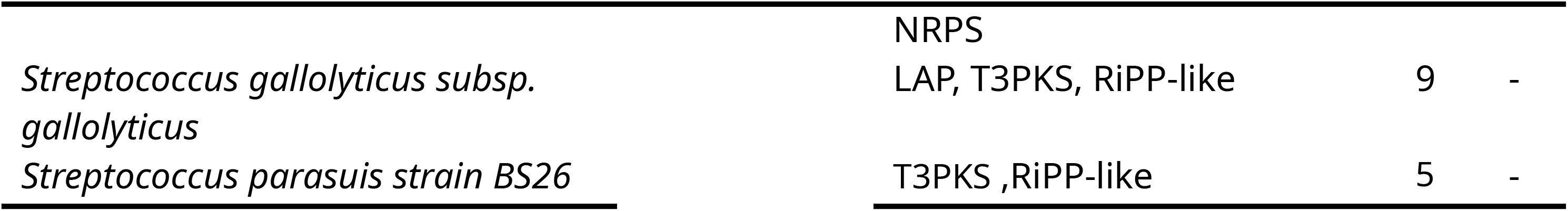
Biosynthetic gene clusters present in genomes from pathogenic and non-pathogenic strains.

**Figure S1.**
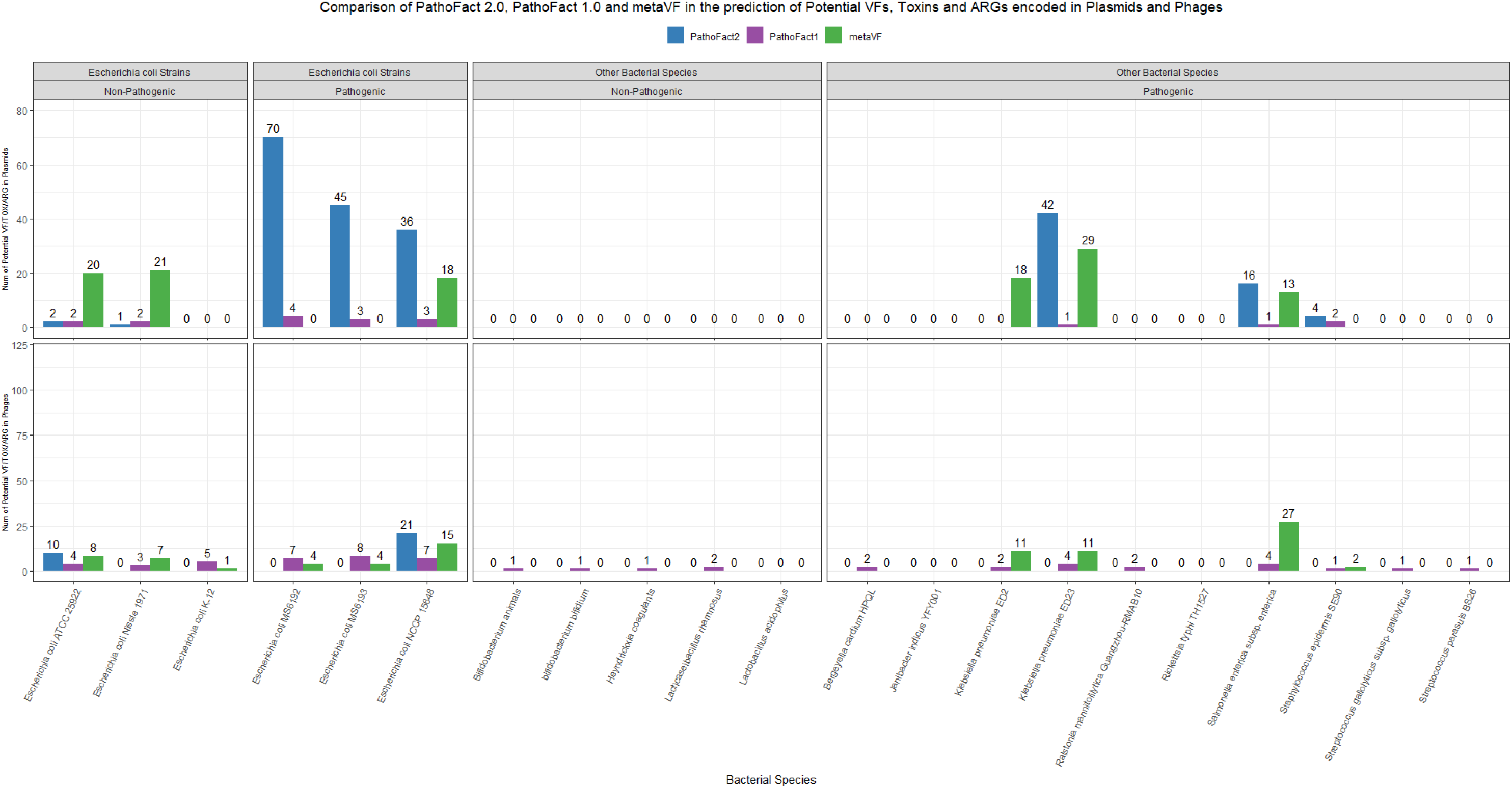
Comparison of PathoFact 2.0 (in blue), PathoFact 1.0 (in purple) and metaVF (in green) in the prediction of potential virulence factors (VFs), toxins, and antimicrobial resistance genes (ARGs) encoded in Plasmids and prophages.

